# Dissecting the Genetic Architecture of Carbon Partitioning in Sorghum using Multiscale Phenotypes

**DOI:** 10.1101/2022.02.16.480719

**Authors:** J. Lucas Boatwright, Sirjan Sapkota, Matthew Myers, Neeraj Kumar, Alex Cox, Stephen Kresovich

## Abstract

Carbon partitioning in plants may be viewed as a dynamic process composed of the many interactions between sources and sinks. The accumulation and distribution of fixed carbon is not dictated simply by the sink strength and number but is dependent upon the source, pathways, and interactions of the system. As such, the study of carbon partitioning through perturbations to the system or through focus on individual traits may fail to produce actionable developments or a comprehensive understanding of the mechanisms underlying this complex process. Using the recently published sorghum carbon-partitioning panel, we collected both macroscale phenotypic characteristics such as plant height, above-ground biomass, and dry weight along with microscale compositional traits to deconvolute the carbon-partitioning pathways in this multipurpose crop. Multivariate analyses of traits resulted in the identification of numerous loci associated with several distinct carbon-partitioning traits, which putatively regulate sugar content, manganese homeostasis, and nitrate transportation. Using a multivariate adaptive shrinkage approach, we identified several loci associated with multiple traits suggesting that pleiotropic and/or interactive effects may positively influence multiple carbon-partitioning traits, or these overlaps may represent molecular switches mediating basal carbon allocating or partitioning networks. Conversely, we also identify a carbon tradeoff where reduced lignin content is associated with increased sugar content. The results presented here support previous studies demonstrating the convoluted nature of carbon partitioning in sorghum and emphasize the importance of taking a holistic approach to the study of carbon partitioning by utilizing multiscale phenotypes.

## Introduction

The integration of multi-scale phenotypes and appropriate mathematical models can assist in the identification of cross-scale interactions leading to emergent properties of dynamic biological systems [1, 2]. Indeed, a holistic understanding of complex systems such as plant above-ground biomass and carbon partitioning requires multiscale phenotypes to address changes in anatomical and physiological processes dictated by underlying genetic networks [3]. The responsiveness of plant carbon-partitioning regimes to environmental conditions such as those induced by a changing climate can significantly affect crop yields and food security thus requiring attention both regionally [4–7] and systemically – particularly under conditions of elevated CO2, heat, drought, and other severe-weather events [8–11]. Crops in the Andropogoneae tribe such as maize [*Zea mays* (L.)], miscanthus [*Miscanthus x Giganteus* (Greef et Deuter)], sorghum [*Sorghum bicolor* (L.) Moench], and sugar cane [*Saccharum officinarum* (L.)] have been the focus of continued development to serve as staple and/or energy crops under extreme weather conditions [12–16] and limit ongoing carbon emissions from fossil fuel use [17–22]. These grasses exhibit highly efficient C4 photosynthetic pathways [23, 24], leaf-level nitrogen-use efficiency [25, 26], water-use efficiency [13, 14, 27], and high yields [26, 28].

Sorghum, in particular, is capable of rapidly accumulating significant quantities of carbon and has been designated as an advanced biofuel feedstock by the U.S. Department of Energy. The Code of Federal Regulations (7 C. F. R. §4288.102) states that advanced biofuels may be derived from biomass in the form of cellulose, hemicellulose, or lignin as well as from sugar or starch [29]. Sorghum meets these conditions as it exhibits great diversity in these carbon-partitioning regimes [30], and the sorghum types are further classified based on these traits as cellulosic, forage, grain, or sweet [29]. Sorghum not only meets the requirements as an advanced biofuel feedstock but is capable of rapidly accumulating significant quantities of non-structural [31] and structural carbohydrates [22, 32, 33] necessary for biofuel [28], forage [34], and grain production [35]. As such, sorghum represents an excellent system for the study of carbon accumulation, partitioning, and designing [29].

Sucrose is the primary source of energy and carbon in plant sink tissues [36] as well as the primary target for ethanol-based, renewable biofuel production [28, 37]. Synthesis of sucrose occurs in the leaf cytosol after which it is transported to various sinks including both storage sinks (i.e., stems) in addition to structural vegetative and reproductive organs, which function as growth sinks [37–39]. However, changes in the quantities of structural and non-structural carbohydrates do not occur in a one-to-one manner nor are they independent [40, 41]. Reduced shoot biomass associated with *dw3* has been shown to decrease grain yield via reduced grain size [42], and differences in carbon partitioning in the stem contribute to tradeoffs between structural and non-structural carbohydrate content [31]. Carbon partitioning is also subject to environmental conditions such as those that transition plants between growth and reproductive phases as seen under drought conditions [13]. A comprehensive examination of the carbon partitioning sinks is necessary to understand the correlations and tradeoffs between these traits in the form of macroscale phenotyping of traits such as above-ground biomass and plant height to the microscale assessment of compositional traits using tools such as near infrared spectroscopy (NIR) [33, 37, 43].

The sorghum Carbon-Partitioning nested association mapping (CP-NAM) panel [29] contains 11 subpopulations generated using diverse parental accessions from the sorghum Bioenergy Association Panel (BAP) [33] and the recurrent parent, Grassl – an accession capable of accumulating significant biomass and fermentable carbohydrates per unit time and area [44]. NAM populations contain sets of RIL families generated from the diverse founders, and as such, benefit from recombination of the founder genotypes, high allele richness, higher statistical power, and are less sensitive to genetic heterogeneity [29, 45]. As the CP-NAM covers the diversity of sorghum types and carbon-partitioning regimes, it represents an excellent source of genotypic and phenotypic diversity to elucidate the genetic architecture underlying carbon fixation, translocation, and utilization so that source/sink dynamics and compositional traits may be understood holistically while simultaneously meeting the demands dictated by a changing environment [29]. Here, we employ quantitative trait locus (QTL) mapping, univariate linear-mixed models (LMMs), and multivariate-response linear-mixed models (MV-LMMs) to identify loci associated with the primary carbon sinks represented by structural and non-structural carbohydrate content in sorghum. Using publicly available genomic resources from the sorghum CP-NAM [29], we identify numerous putative loci involved in carbon partitioning both known and novel.

## Materials and Methods

### Plant Materials and Phenotyping

CP-NAM seeds were accessed from the Clemson University sorghum germplasm collection and planted at the Simpson Research Farm (34.64737384683981, −82.74780269784793), South Carolina in May of 2020. While the original CP-NAM panel contained 2,489 accessions, a subset of 110 individuals were selected from each RIL family using the Partitioning Around Medoids (PAM) function in the R package *cluster* [46] to reduce the field size and manual labor necessary to manage the CP-NAM field while representing most of the genetic diversity within the RIL populations [47]. A total of 110 sample clusters were identified based on the genomic data from each RIL family with a medoid sample centrally located in each cluster (Figure 1). The medoid is the individual within the cluster that best represents the genetic diversity of that sample cluster. The 110 accessions representing the medoids for each population were selected as representatives of each cluster and planted for phenotyping in 2020.

**Figure 1.**
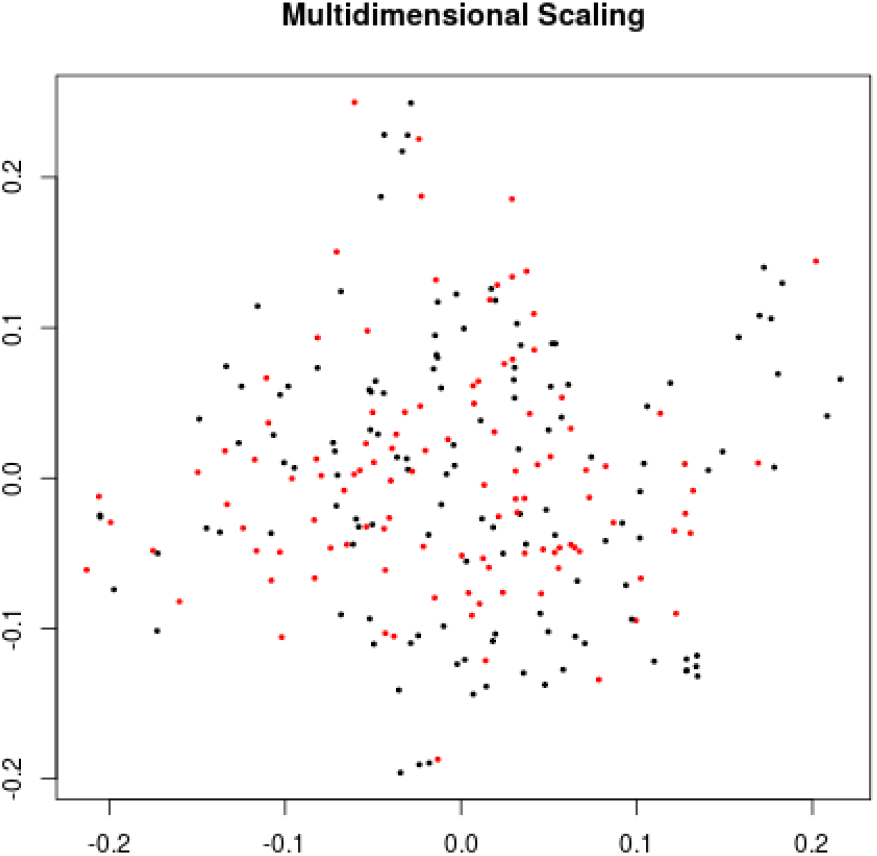
A multidimensional scaling plot using the genotypic data of PI229841 RIL accessions where the 110 medoid samples are colored red and the remaining samples are black. The x-axis represents the first principal component of the genotypic data, and the y-axis represents the second principal component. The proximity of accessions to each other indicates the degree of similarity – with shorter distances indicating higher similarity.

The NAM RILs were planted in single-row plots that were 3.04 m in length with 0.762 m between-row spacing in a randomized complete block design with two replicates per line. Randomization was done within blocks containing RILs from a given family, and families were planted together to avoid large field effects within families. Plants were irrigated on an as-needed basis but did not occur past 90 days after planting due to plants exceeding the irrigation pivot height. Harvesting started September 14th and continued through October 19th. Due to the range in harvesting days, the maturity stage and days to harvest were recorded for each plot to use as covariates for phenotypic and genomic analysis to avoid the confounding effect of varying maturity groups across the RIL families. Phenotypes collected included above-ground biomass, stand count, maturity at harvest, days to harvest (DTH), plant height, and dry and wet weights. Stand count was measured as the total number of emerged seedlings between 15-30 days after planting. Plant height was measured at harvest from the base of the stalk to the apex of the panicle, or, if no panicle was present, to the apex of the shoot apical meristem.

A representative meter was selected for each plot, and all stalks were cut at the base within that meter and weighed (in kilograms). From those stalks, three representatives were selected for subsequent phenotyping including above-ground biomass (including panicles and leaves), wet weight, and dry weight, where biomass represents a scaled meter weight. Any panicles or partially formed panicles were removed along with all leaf matter before collecting wet weight. The stalks were then cut into billets for collection into mesh drying bags and placed into drying bins at 40 °C until stalks were dried to a constant moisture content. Dried stalks were removed from the drying bins, and dry weights were taken. Stalks were then ground with a Retsch SM 300 cutting mill so that compositional traits could be measured using a PerkinElmer DA7250™ NIR instrument (https://www.perkinelmer.com), which uses calibration curves for spectral measurements built using wet chemistry values generated by Dairyland Laboratories, Inc. (Arcadia, WI, USA) as described in (Brenton2016-lf). Compositional traits include acid detergent fiber (ADF), adjusted crude protein (Adj. CP), neutral detergent fiber (NDF), ash-free NDF (aNDFom), ash, calcium, chloride, dietary cation-anion difference (DCAD), dry matter, potassium, lignin, magnesium, moisture, net energy growth (NEG), net energy lactation (NEL), relative feed value (RFV), non-fiber carbohydrates (NFC), and water-soluble carbohydrates (WSC) where all traits are expressed as a percent of dry matter. NEG, NEL, NEM, and TDN were also estimated using an Ohio Agricultural Research & Development Center (OARDC) summative energy equation and may appear conjugated with the OARDC abbreviation (Figure 2).

**Figure 2.**
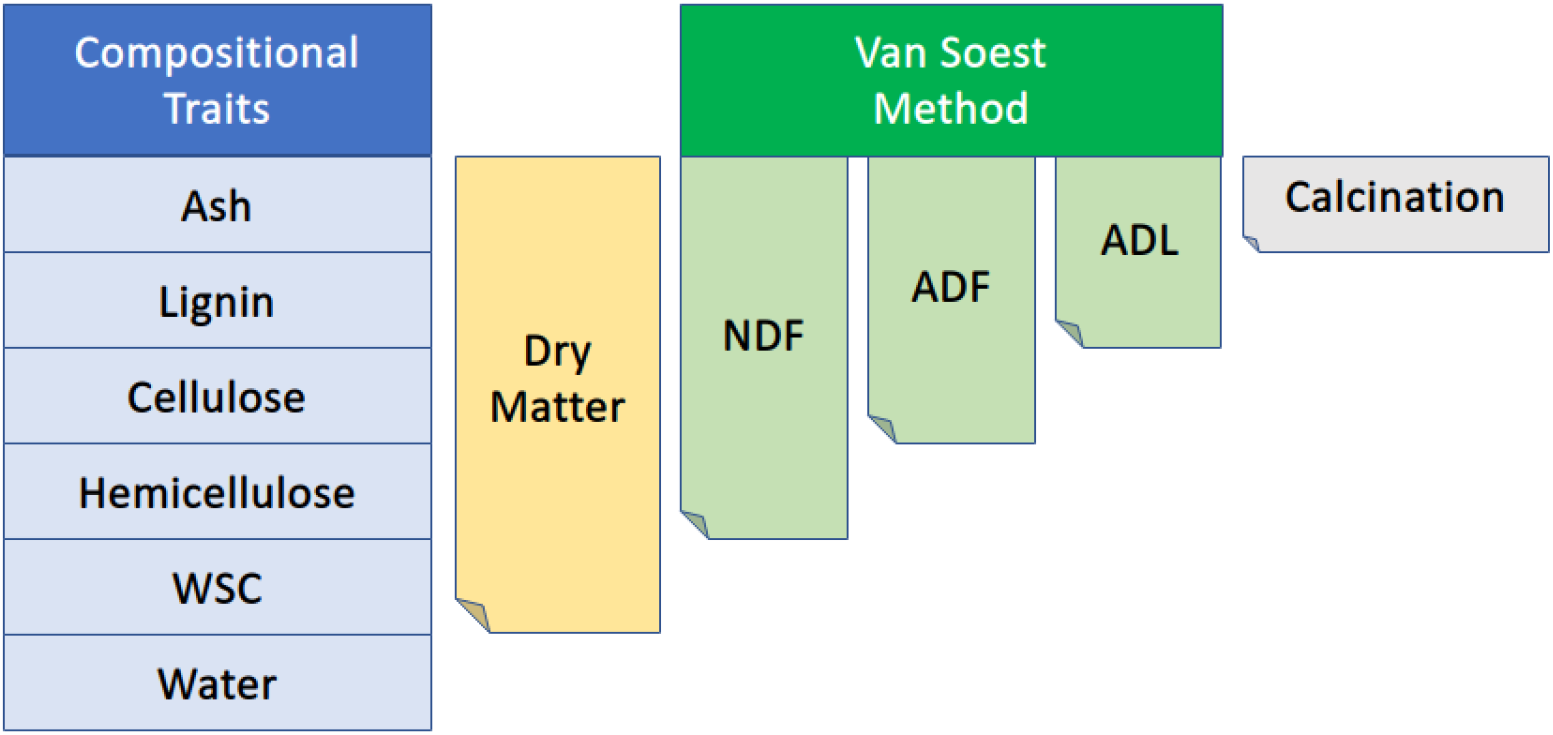
Diagram of the relationship among several compositional traits collected via NIR. NDF, neutral detergent fiber; ADF, acid detergent fiber; ADL, acid detergent lignin; WSC, water-soluble carbohydrates. The Van Soest method permits the distinction of soluble cell contents from the less digestible cellular components such as lignin, cellulose, and hemicellulose. Figure adapted from [48].

### Descriptive Statistics and QTL Mapping

The repeatability was estimated for all traits using the package *heritability* in the R programming language (R Core Team 2019) [49]. The best linear unbiased predictors (BLUPs) were calculated for each trait with the R package *lme4* [50] using the *lmer* function with genotype as random effects. The resulting BLUPs were used as adjusted phenotypic values for all mapping and association analyses. Heatmaps and correlation metrics were measured using the *seaborn* [51] and *pandas* [52] packages in CPython [53], respectively.

Genotypic data from [29] were used to perform QTL mapping and genome-wide association studies (GWAS). In summary, Genotyping-By-Sequencing (GBS) data were generated using a double-digest approach (*PstI* and *MspI*), processed using the Tassel GBS version 2 pipeline [54], and imputed using Beagle 5.1 [55] as described in [29]. Genetic maps were constructed for each RIL family with Haldane’s mapping function [56] and a genotyping error rate of 0.0001, where the conditional probabilities of the true genotypes were estimated using a hidden Markov model. Map construction and QTL mapping were performed for each RIL family using the *qtl2* package [57] in R, and genomic scans were performed using Haley-Knott regression [58] and linear mixed model approaches [57] including both full and leave-one-chromosome-out models. All models included maturity and DTH as covariates except for DTH. QTL effects were estimated as 100 × (1 – 10^(−2×*LOD*)/*n*^), where *n* is the number of individuals in the corresponding mapping population [59]. However, we recognize that estimates for PVE with these population sizes may exhibit inflated values due to the Beavis effect [60].

### Genome-Wide Association

GWAS were done using both the Genome-wide Efficient Mixed Model Association (GEMMA) program version 0.98.3 [61] and the Genome Association and Prediction Integrated Tool (GAPIT) version 3 [62, 63]. We specifically used GEMMA for both MV-LMMs and Bayesian Sparse Linear Mixed Models (BSLMM) [64] while GAPIT was used for BLINK [65] and LMMs. The phenotypic and genotypic data were converted to Plink format using Plink [v1.90b6.10] [66]. The genomic relatedness matrix was calculated using the VanRaden algorithm [67] and all models were run using a MAF filter of 0.01. Univariate LMMs are of the following the form,

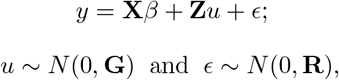

where **y** is a vector of phenotypic values for a single trait, **X** is the numeric genotype matrix generated from the SNPs, *β* represents the unknown vector of fixed effects representing the effect size for each SNPs, **Z** is the design matrix for random effects, **u** is the unknown vector of random effects, and *ϵ* is the unknown vector of residuals. These univariate models test the alternative hypothesis *H*_1_: *β* ≠ 0 against the null hypothesis *H*_0_: *β* = 0 for each SNP.

In addition to the frequentist univariate model described above which estimates fixed effect coefficients by selecting the optimal value minimizing the least-squared error – the equivalent of a flat prior, we ran a BSLMM which assumes fixed effects are distributed according to the sparse prior, *β ~ πN* (0, *σ*2*aτ* – 1) + (1 – *π*)*δ*_0_ [64]. Runs were executed using 20e6 sampling steps with a burn-in of 5e6, and the posterior inclusion probability (PIP) threshold established as 0.036 based upon a 99.95% quantile from simulated data sets across quantiles to determine the empirical significance cutoff [68]. While more computationally intensive due to the Markov Chain Monte Carlo sampling approach involved, this model provides shrinkage of *β* estimates to control for type I errors and provides a posterior distribution of plausible values rather than simple point estimates. Additionally, we also conducted univariate analyses using BLINK [65]. BLINK approximates the maximum likelihood approach used by LMMs, instead using Bayesian Information Criteria in a fixed effect model where each SNP is iteratively associated with the phenotype of interest. Markers in linkage disequilibrium (LD) with the most significant marker are then excluded – as estimated using a Pearson correlation coefficient ≤ 0.7. For subsequent markers, the next most significant SNP is selected, and the exclusion process is conducted in the same way until no markers can be excluded. Unlike SUPER and FarmCPU methods, BLINK does not assume that causal genes are evenly distributed across the genome and is faster with higher statistical power – due to its multi-locus approach – and lower type I error rates [65].

For MV-LMMs, we used GEMMA models of the form:

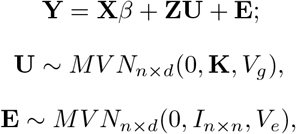

where **Y** is an n by d matrix of d phenotypes for n individuals, **X** is the numeric genotype matrix generated from the SNPs, *β* is a d vector of fixed effects representing the effect size for the d phenotypes, **Z** is the design matrix for random effects, **U** is the n by d matrix of random effects, **E** is the n by d matrix of residuals, **K** is a known n by n relatedness matrix, *V*_*g*_ is a d by d symmetric matrix of genetic variance components, *I*_*n*×*n*_ is an n by n identity matrix, and *V*_*e*_ is a d by d symmetric matrix of environmental variance components. As the maturity of accessions significantly affects all phenotypes, maturity and DTH were used as covariates in all QTL mapping and GWAS models except for DTH.

### Multivariate Adaptive Shrinkage

The estimated effect sizes and standard errors for every marker in the the LMMs for ADF, ash, dry matter, dry weight, height, NDF, P, wet weight, and WSC were filtered using a local false sign rate < 0.1 based on a condition-by-condition analysis using ashr in R [69]. A control set of estimated effects and standard errors were also randomly selected for 40,000 markers to estimate the covariance between SNPs for each phenotype. A correlation matrix of the random control set was estimated and used to control for the confounding effects of correlated variation among the traits using mashr in R [70]. We utilized both canonical and data-driven covariance matrices following mashr best practices to test for pleiotropy across traits [70]. The posterior probabilities were calculated for each SNP by fitting a mash model with all tests. Bayes factors were extracted and plotted from mash results using the CDBNgenomics R package [71]. Variants exhibiting Bayes Factors greater than 10 were considered as demonstrating significant pleiotropic effects.

## Results

### Trait Heritabilities and Correlations

Heritability is the proportion of phenotypic variance attributable to genetic variance, and when differences between genotypes is assumed to derive entirely from genetic effects, the measurement of consistent individual differences is called repeatability [49]. We calculated the repeatability for all traits and identified many traits with repeatability greater than 0.2 (Table 1) with maturity and DTH exhibiting the highest repeatabilities (> 0.75). Agronomic phenotypes exhibited higher repeatability compared to compositional traits with all agronomic traits exceeding 0.5 repeatability. Agronomic traits also demonstrated relatively low correlation among the traits except for biomass, wet weight, and dry weight which were all highly correlated (≥ 0.67) (Figure 3). These measures for repeatability and correlation among traits are consistent with previous estimates in sorghum [33].

**Table 1.** Repeatability for agronomic (top portion of table) and compositional (bottom portion) traits. Agronomic traits include maturity at harvest, plant height, wet weight, dry weight, above-ground biomass, and days to harvest. Compositional traits include acid detergent fiber (ADF), adjusted crude protein (Adj. CP), ash-free neutral detergent fiber (aNDFom), crude protein, dry matter, lignin, neutral detergent fiber (NDF), net energy growth (NEG), total digestible nutrients (TDN) as a percentage of ADF, and water-soluble carbohydrates (WSC). NEG was estimated using an Ohio Agricultural Research & Development Center (OARDC) summative energy equation.

| Trait | Repeatability |
| --- | --- |
| Maturity | 0.81 |
| Plant Height | 0.53 |
| Wet Weight | 0.52 |
| Dry Weight | 0.55 |
| Biomass | 0.52 |
| DTH | 0.76 |
| ADF | 0.65 |
| Adj. CP | 0.38 |
| aNDFom | 0.61 |
| Crude protein | 0.35 |
| Dry Matter | 0.36 |
| Lignin | 0.45 |
| NDF | 0.55 |
| NEG (OARDC) | 0.59 |
| WSC | 0.47 |
| TDN (ADF) | 0.62 |

**Figure 3.**
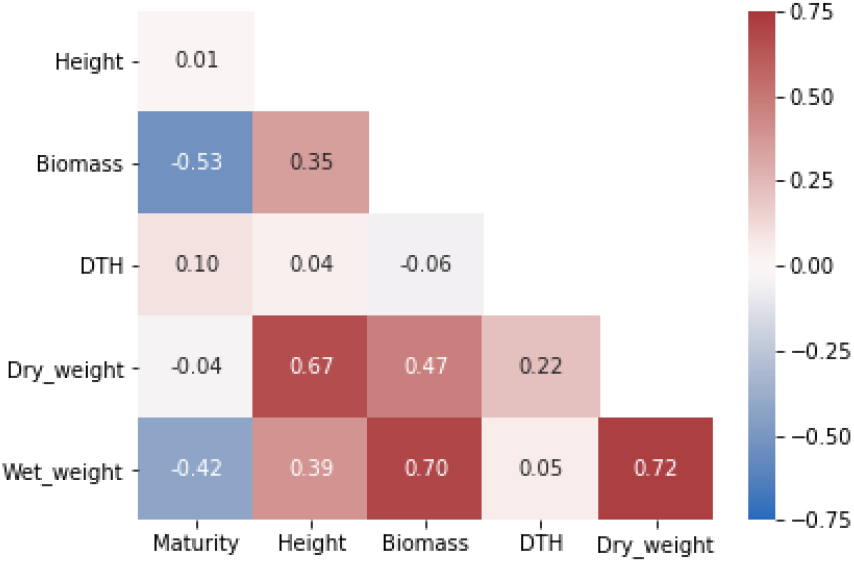
Heatmap of Pearson’s correlation coefficients among agronomic phenotypes. Biomass represents a scaled meter weight, and DTH is days to harvest.

Many compositional traits exhibited strong correlation (> |0.5|) (Figure 4), which is expected due to the aggregate nature of some traits and their dependency on maturity (Figure 2). Importantly, while many of the fiber-based compositional traits exhibited strong repeatability, only six compositional traits of 34 had Pearson’s correlation coefficients > |0.3| with dry weight and none had values exceeding |0.5.| The lack of strong correlation between compositional traits and dry weight suggests that sorghum composition could be improved without significantly affecting total vegetative yield [33, 43].

**Figure 4.**
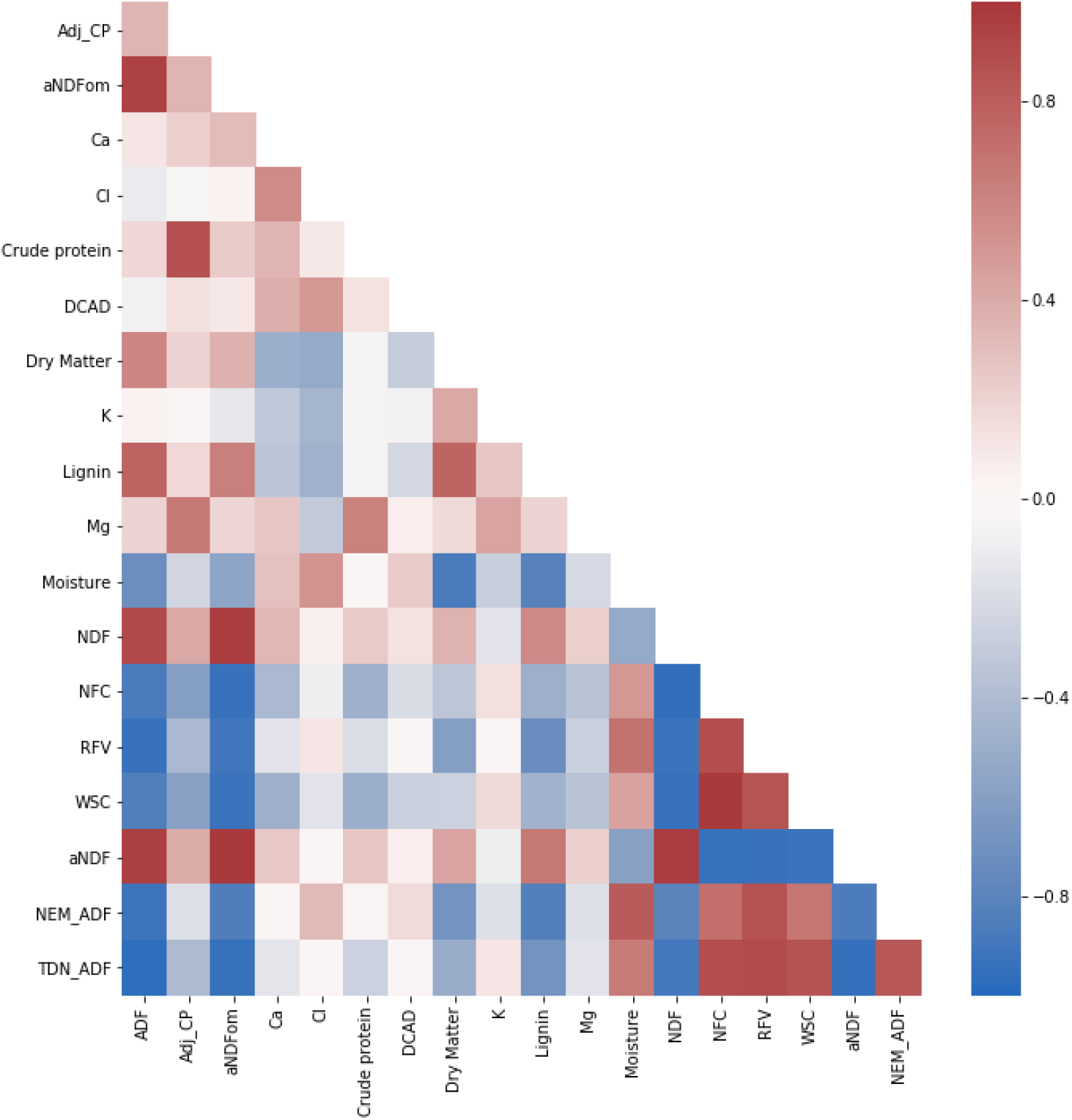
Heatmap of Pearson’s correlation coefficient among compositional traits. Traits include acid detergent fiber (ADF), adjusted crude protein (Adj.CP), neutral detergent fiber (NDF), ash-free NDF (aNDFom), calcium, chloride, crude protein, dietary cation-anion difference (DCAD), dry matter, potassium, lignin, magnesium, moisture, non-fiber carbohydrates (NFC), relative feed value (RFV), and water-soluble carbohydrates (WSC). NEG, NEL, NEM, and TDN were also estimated using an Ohio Agricultural Research & Development Center (OARDC) summative energy equation and may appear conjugated with the OARDC abbreviation.

## Mapping and Associations

### Agronomic Traits

As the CP-NAM is composed of 11 RIL families, it provides the opportunity to resolve genotype-to-phenotype associations through both QTL mapping and GWAS. To this end, QTL mapping was performed for each RIL family in the CP-NAM for every trait using maturity and DTH as covariates except when DTH is the response variable. Several known QTL were identified for plant height on chromosomes (Chr) 6 [*qHT7.1* /*Dw2*], Chr7 [*Dw3*], and Chr9 [*Dw1*] aggregated by RIL families derived from grain, cellulosic, sweet, and forage parents (Figure 5; Table S2) along with several potentially novel associations on Chr1 and Chr8. The newly identified QTL on Chr1 spanned from 10 to 12 Mb (13.2 PVE) and from 22 to 56.7 Mb (14.5 PVE) (Table QtlAgronomic). The QTL from the latter position also had a significant genome-wide association for height using the BLINK model with a peak at Chr1:50,888,855 (Figure S1). Another novel QTL was identified for height on Chr8 from approximately 0.37 to 2.7 Mb (Table Chr8Height) using a leave-one-chromosome-out method (Figure 5c; Figure S2). A significant genome-wide association for height was found for the SNP at Chr8:2,033,695 using the BSLMM model (Figure S3), and the associated region is within previously identified QTL for transpiration rate and efficiency of energy of PSII [72].

**Figure 5.**
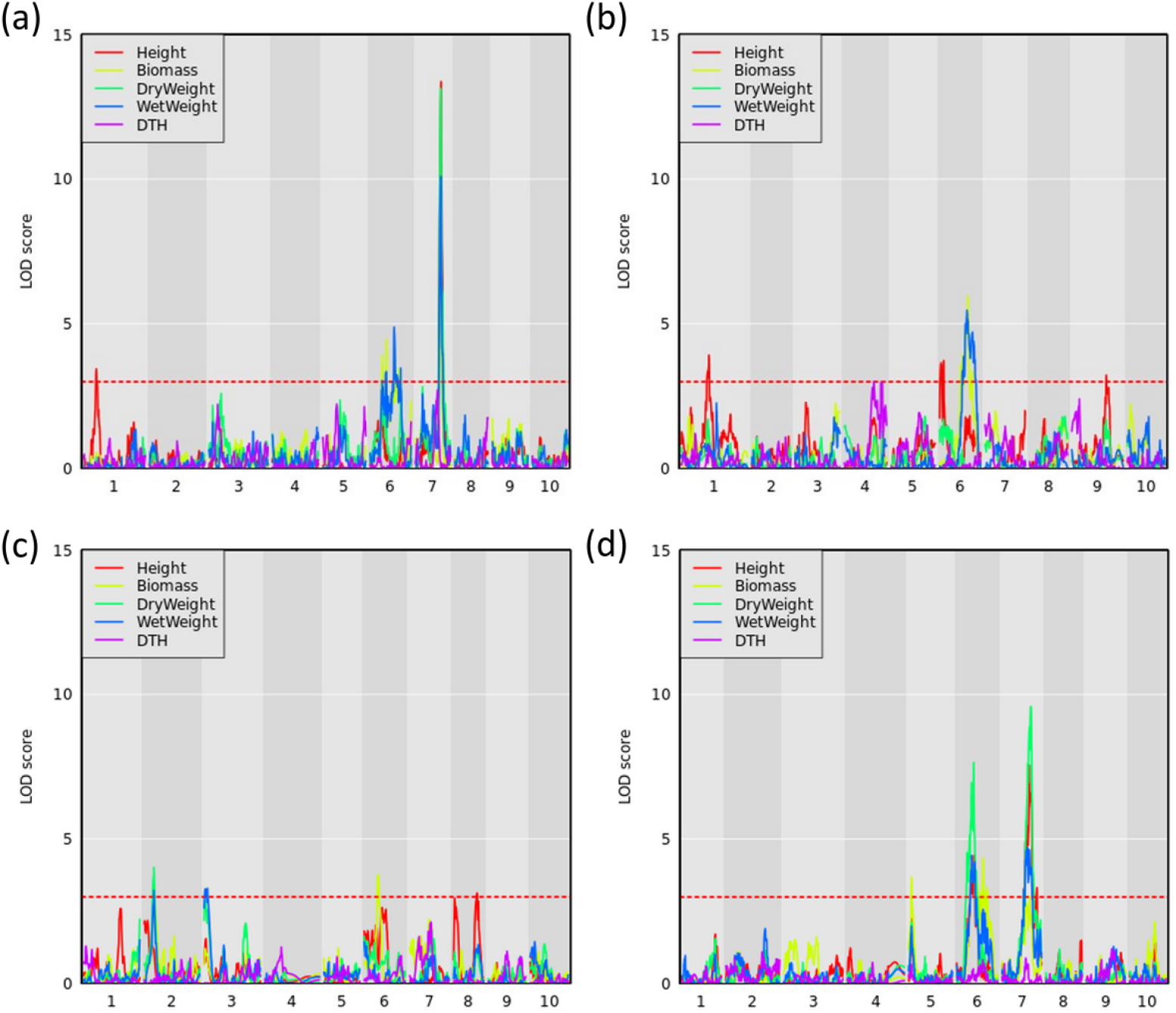
QTL mapping for agronomic traits with maturity covariate where the red dashed line represents a logarithm of the odds (LOD) threshold of three in (a) PI297155, (b) PI506069, (c) PI586454, and (d) PI655972 RILs, which are grain, cellulosic, sweet, and forage recombinant populations, respectively.

Sorghum has over 40 identified flowering time and maturity QTL [73]. QTL mapping results for DTH in PI506069 RILs identified a locus on Chr4 (11.8 PVE) from 70 to 113 cM that peaked at 79 cM (Figure 5b). This QTL colocalizes with the flowering time gene *CN2* [74], which is a centroradialis-like gene homologous to Terminal Flower1 (*TFL1*). An additional 11 loci were identified using BLINK including *Ma3* /*Ma5* [Chr1], *SbCN12* [Chr3], and *Ma1* [Chr6] [75] along with several other unidentified loci (Figure S4). The identified loci include phytochromes and other flowering time modulators that mediate photoperiod sensitivity in these non-temperately adapted accessions.

### Biomass Traits

In addition to height and DTH phenotypes, various measures of biomass yield were taken including wet weight, dry weight, and above-ground biomass (abbreviated as biomass). These biomass traits were often associated with the same QTL – particularly the QTL on Chr6 and Chr7 (Figure 5), but significant associations from GWAS were more variable (Figure 6). The QTL on Chr3 (12.7 PVE) identified using wet weight spans from approximately 1 to 6 Mb in the sweet x cellulosic RILs of PI586454 (Figure 5c) and coincides with QTL associated with stem circumference and transpiration rate [72, 76]. GWAS of wet weight also identified an association on Chr3 at approximately 62Mb (Figure 6d), which colocalizes with numerous trait associations including plant height [77], stem circumference [76], and days to flowering [78]. However, the gene(s) mediating these phenotypes is unclear. Dry weight and wet weight were associated with several QTL on Chr6 (Figure 5a, d) that were also captured through GWAS (Figure 6) and ranged from 1 to 5 Mb and 49 to 51 Mb, respectively. The QTL spanning 1 to 5 Mb corresponds to the known maturity locus, *Ma6* [79]. These phenotypes also captured the height loci *Dw2*, which encodes a protein kinase that regulates stem internode length [80], and *Dw3*, which encode a P-glycoprotein auxin transporter and only affects plant height below the flag leaf [81]. As auxin stimulates the production of hemicellulose and consequently stem elongation, the association is consistent with the known identity of the locus [81].

**Figure 6.**
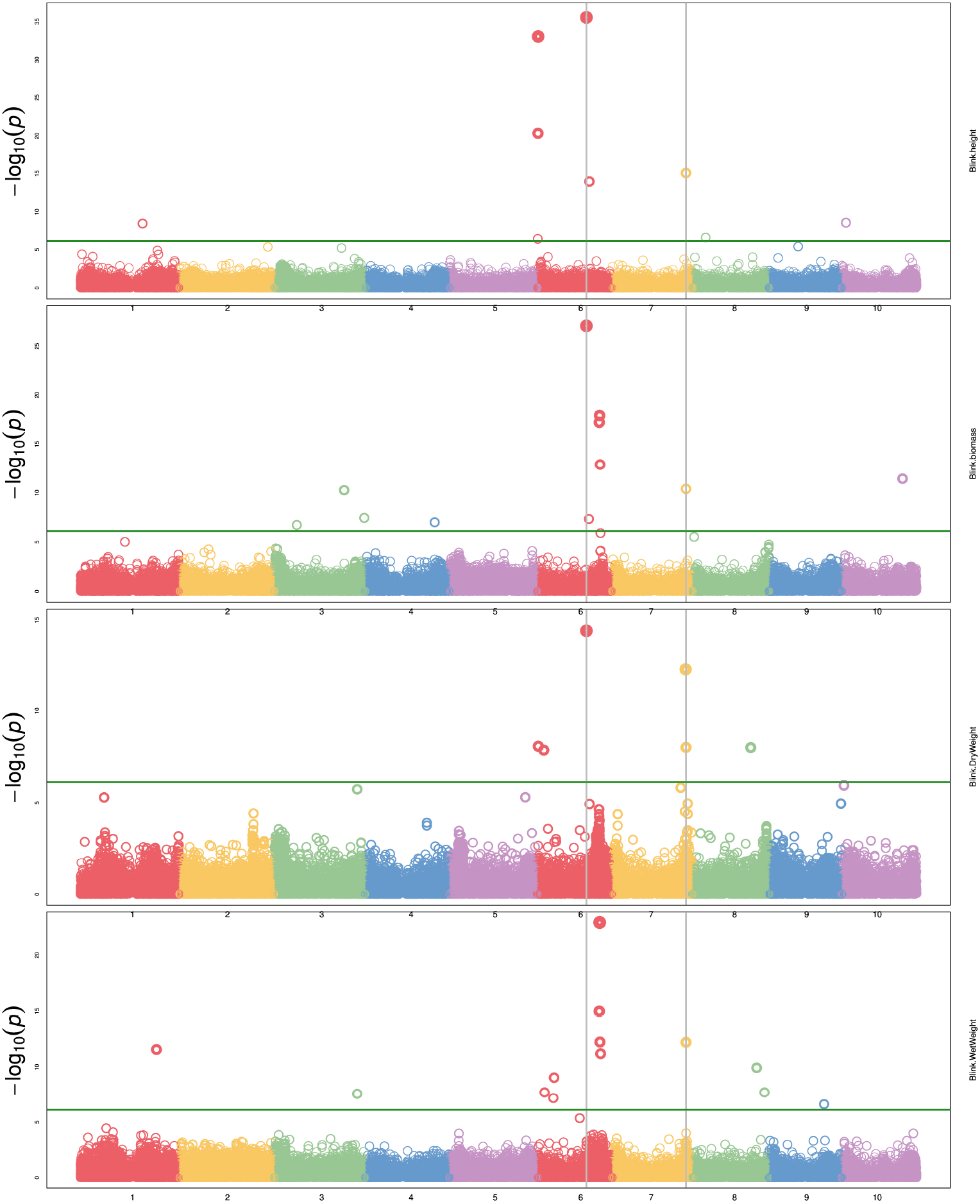
Manhattan plot of several agronomic traits using BLINK with maturity and DTH as covariates. (a) plant height, (b) biomass, (c) dry weight, and (d) wet weight. The green horizontal line represents the Bonferroni-corrected significance threshold. Vertical lines indicate SNPs common to multiple phenotypes.

The forage RIL family, CP-NAM PI655972, was uniquely suited for identifying a QTL (14.3 PVE) controlling biomass content on chromosome 5 (Figure 5d). The biomass QTL also overlapped a QTL for adjusted crude protein content (Table QTLCompositional). [82] identified a sucrose content QTL that falls completely within the biomass QTL and partially overlaps the adjusted crude protein QTL seen here [82]. Given the large range of the QTL or even the overlapping region, it is difficult to pin down what gene(s) may be responsible for these associations. In addition to this unique locus, biomass was associated with the same loci on Chr6 and Chr7 as height, wet weight, and dry weight (Figure 6).

### Compositional Traits

QTL mapping was also performed for all compositional traits, and select traits were plotted for all RIL families (Figure 7). Several overlapping QTL were identified across traits within RIL families, and the various sorghum RIL families/types captured different QTL for the same traits. The most significant QTL on Chr6 associated with ADF was consistently identified in all RIL families. The narrowest range of this locus was obtained in PI229841 and PI508366 RILs and spanned from approximately 50.3 to 51.8 Mb (Table QtlNIRTable).

**Figure 7.**
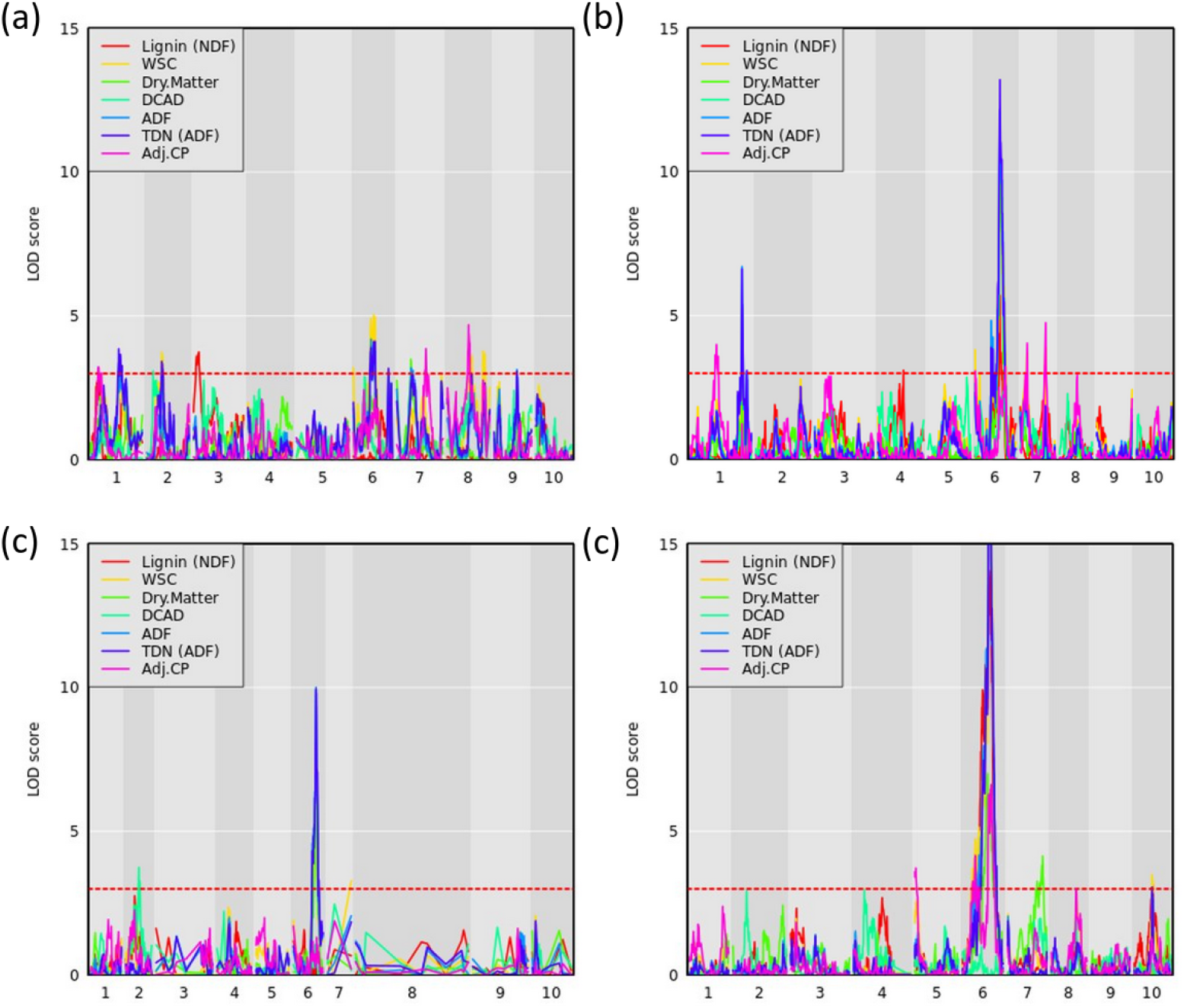
QTL mapping for compositional traits with maturity and DTH covariates where the red dashed line represents a logarithm of the odds (LOD) threshold of three in (a) PI22913, (b) PI297155, (c) PI508366, and (d) PI655972 RILs, which are sweet, grain, cellulosic, and forage recombinant populations, respectively.

WSC content provides an estimate of the carbon partitioned and accumulated by accessions in the stem in the form of water-soluble carbohydrates [37]. For WSC, we identified a QTL on Chr6 in the PI22913 RILs around 50 Mb, which results from a cross between the sweet sorghum accession PI22913 and the cellulosic Grassl (Figure 7a). The QTL occurs within the Dry Midrib (*D*) locus, which has also shown strong association with midrib color, grain yield, sugar yield, juice volume, and biomass, indicative of a pleiotropic effect of the *D* locus across these phenotypes [83, 84]. This QTL also overlaps with QTL identified using ADF, wet weight and dry weight phenotypes described above. [83] previously demonstrated that green midrib color was more strongly associated with sugar content traits than the *D* locus genotypic data with sugar content, and therefore suggested that selecting for green midribs is a simple alternative to genetic selection for sweet sorghum breeding programs. Consistent with this observation, the *D* locus accounted for 64.2% of the variance explained for WSC in PI229841 RILs.

Following the design pattern indicated in Figure 2, we used a combination of LMMs with various compositional traits as covariates to deconvolute the contribution of individual traits to phenotypic variance, and as a converse approach, we also ran multivariate-response models on constituent parts to compare with composite traits. While population genomic studies have historically utilized univariate LMMs, more recent works are finding that multivariate-response linear mixed models (MV-LMM) have higher true-positive rates particularly when correlated traits with low, medium, and high heritabilities are analyzed together in one MV-LMM [85]. The use of MV-LMMs may also provide additional power to detect causal loci exhibiting pleiotropic effects across multiple traits [86]. By using MV-LMMs on combinations of carbon-partitioning traits, we can better understand the interplay among these traits and predict the systemic effects of trait selection on the respective carbon sinks.

Running WSC and NDF in a multivariate-response model with maturity and DTH as covariates approximates the dry matter phenotypic variance (Figure 2; Figure 8). While only one locus has significant associations, the highly significant association on Chr6 occurs broadly from approximately 49.5 to 52Mb with the three most significant SNPs (50,556,927; 50,558,124; and 50,574,062). As noted from the QTL mapping results for WSC, this associated corresponds to the D locus. The most significant SNP (Chr6:50,558,124) exhibited considerable phenotypic variation across traits but contrasting effects for NDF and WSC for each allele (Figure S5-S6). The identity of the gene underlying this locus is believed to be a NAC transcription factor where recessive parents possess a premature stop codon in the NAC domain and were shown to exhibit lower lignin content but higher sugar and grain yields [84] similar to the relationship between NDF and WSC seen here. Running the same model but adding the top SNP as a covariate brings the peak on Chr4 above the significance threshold (Figures S7-S8), which is consistent with the dry matter LMM on Chr4 corresponding (Figure 10b). This SNP overlaps a QTL previously identified with dry matter growth rate, leaf appearance rate, [87], and stem circumference [76]. Potential candidate genes in the region include two high-affinity nitrate transporter (NRT) genes (Sobic.004G009400/Sobic.004G009500). In sorghum, increased expression of NRTs has been suggested to improve the efficiency at which inorganic and organic nitrogen is assimilated [88] and affect both the biomass and grain yield [89].

**Figure 8.**
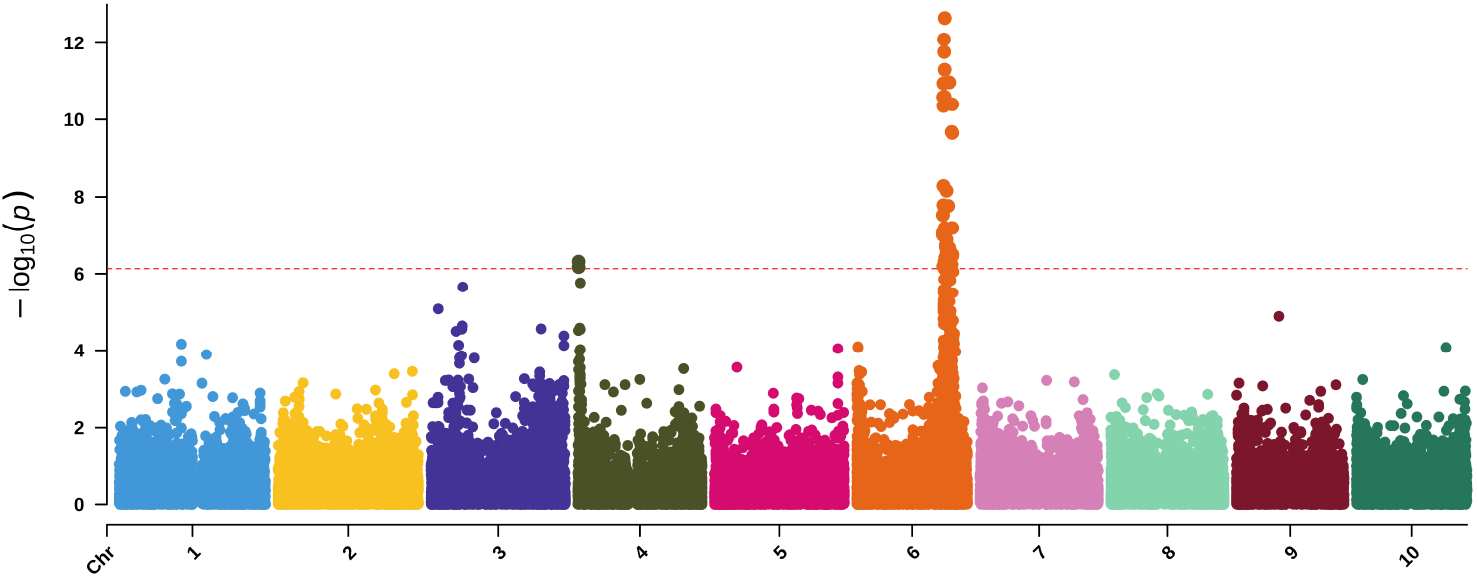
Manhattan plot of a MV-LMM using GEMMA with WSC and NDF as response variables and both maturity and DTH as covariates. The red-dashed, horizontal line represents the Bonferroni-corrected significance threshold.

**Figure 9.**
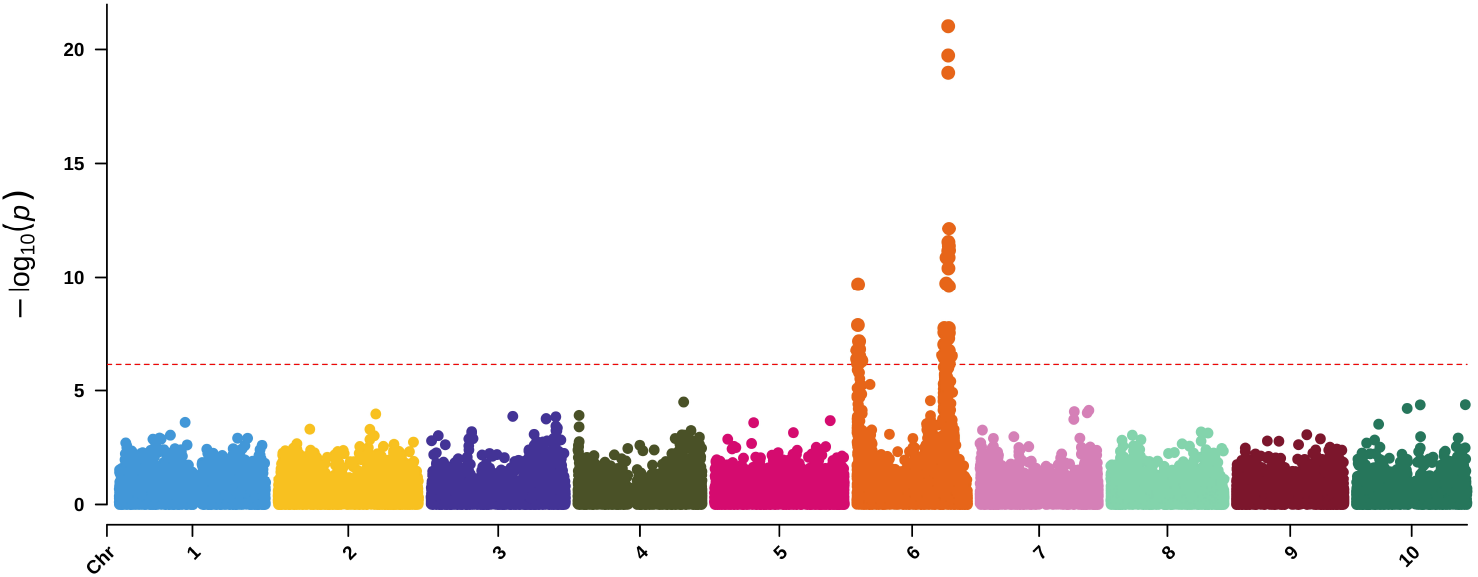
Manhattan plot of a MV-LMM using GEMMA with ash and lignin as response variables and both maturity and DTH as covariates. The red-dashed, horizontal line represents the Bonferroni-corrected significance threshold.

**Figure 10.**
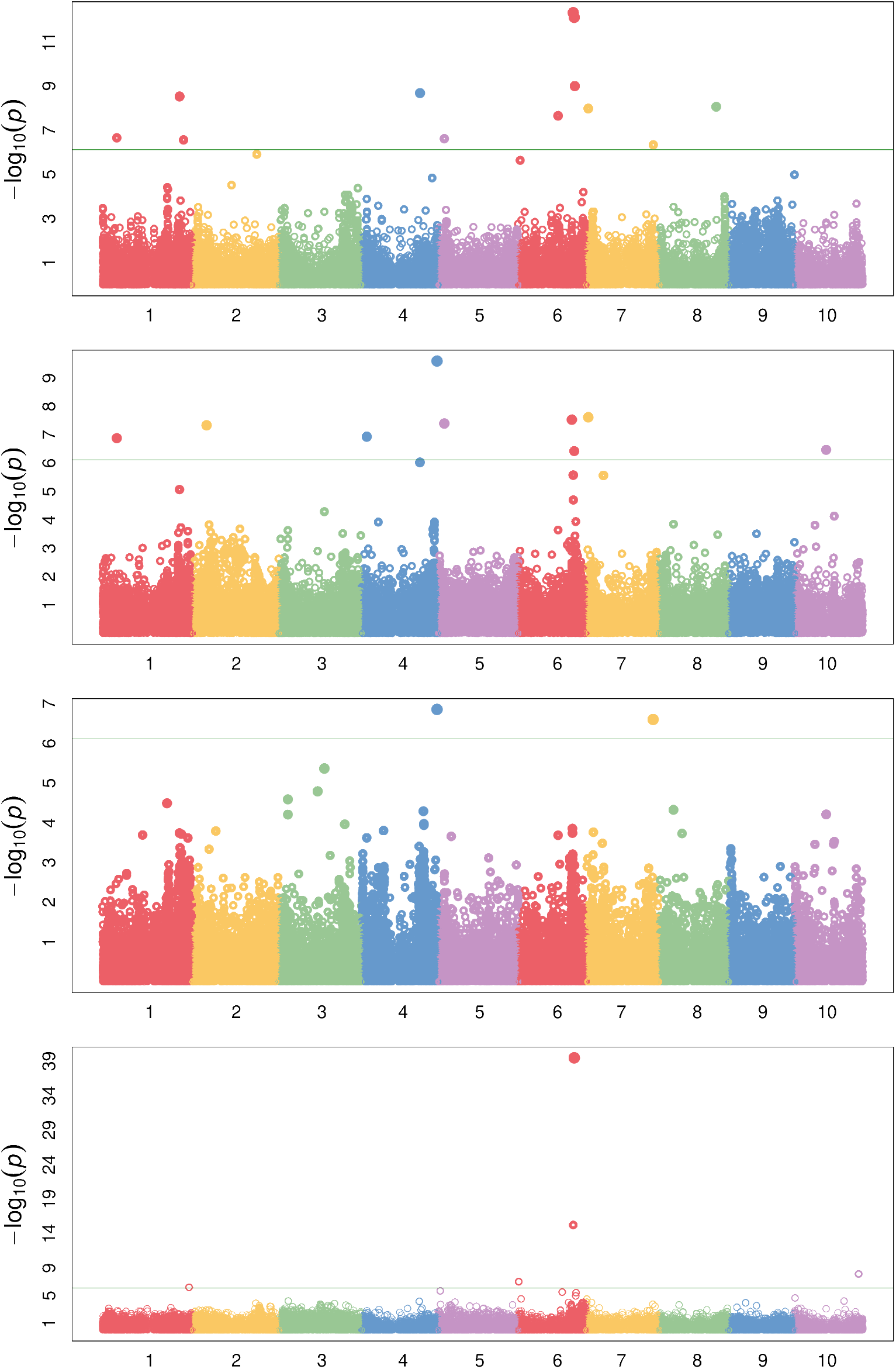
Manhattan plot of several compositional traits with different covariates following the design represented by Figure 1. (a) dry matter with maturity and DTH covariates, (b) dry matter with maturity, DTH, and NDF covariates, (c) dry matter with maturity, DTH, and ADF covariates, and (d) WSC with maturity and DTH covariates. The green horizontal line represents the Bonferroni-corrected significance threshold.

Using MV-LMM, we ran ash and lignin as response variables with maturity and DTH covariates (Figure 9), which may be roughly viewed as examining ADL (Figure 2). Though we do not have a direct measure of ADL for comparison, the significant loci are a subset of those found using ADF and NDF (Figures S9-S10) indicating the utility of MV-LMM and covariate models for compositional analyses.

By examining accumulation of soluble sugars in addition to lignocellulosic content and aggregate traits such as NDF, we may obtain a broader perspective of varying sink strength across the accessions. NDF represents a measure of the total lignocellulosic content. Lower lignin content and high lignocellulose production is preferred for efficient biofuel production [90]. Using dry matter as the response variable, NDF was included in the LMM as a covariate to examine the effects of keeping NDF constant on the SNP significance (Figure 10b). Biologically, this could be seen as running a model on the phenotypic variance of WSC. Similarly, using dry matter as the response variable with ADF as a covariate examines the phenotypic variance of hemicellulose and WSC (Figure 10c). By using covariates or multivariate-response models, the individual (i.e., WSC) or cumulative (i.e., NDF) phenotypic variance of compositional traits may be disentangled, and the relationships among traits may be more clearly distinguished.

Interestingly, the significantly associated SNPs are not identical between models such as a WSC LMM versus a dry matter response with NDF covariate LMM (Figure 10b, d) with composite traits such as dry matter often exhibiting more associated loci. As previously indicated, the integration of multi-scale phenotypes and multivariate models may be identifying emergent properties of these biological systems as some trait associations are not merely the sum of their parts [1, 2], which further indicates the importance of running several different models that attempt to examine characteristics of a trait from multiple perspectives. The various dry matter models (Figure 10a–c; Figure S11) were also associated with a previously identified WSC locus containing a putative vacuolar iron transporter (VIT) on Chr4 [37]. It has been suggested that the candidate gene underlying this locus (Sobic.004G301500) may affect sugar accumulation either through neofunctionalization or via an iron-deficiency response [37]. Though interestingly, the same LD block is also hit with adjusted crude protein as well as NEG (Figure S12-S13). Previous identification of a putative *Dw4* locus identified from plant height GWAS also corresponds to this locus [81], and the locus has also been associated with increased total biomass and root biomass [91]. Together, these results suggest a mechanism foundational to carbon accumulation underlying this locus, or the locus exhibits a pleiotropic effect on carbon accumulation or partitioning.

While the sorghum gene is classified as an iron transporter, further comparison with the Arabidopsis ortholog (AT3G43660) indicates a potential role in cellular manganese ion homeostasis (GO:0030026) [92, 93]. The CCC2-like domain of Sobic.004G301500 or one of the duplicate loci (Sobic.004G301600/Sobic.004G301650) therefore likely acts to transport manganese to vacuoles and maintain manganese homeostasis. As manganese serves to increase nitrogen assimilation [94], is a fundamental catalyst during the water-splitting reaction of photosystem II [95], and is necessary for respiration [96], a pivotal role in manganese homeostasis might better explain these associations. Further, since a role in manganese homeostasis has been described, these duplicated loci may instead demonstrate subfunctionalization followed by tissue-specific expression of one copy or the duplication may alter gene dosage and consequently modify some rate-limiting process. The association on Chr7 at approximately 59.5 Mb found using dry matter with an ADF covariate, which is equivalent to looking at phenotypic variation due to WSC and hemicellulose, is *Dw3* (Figure 10c).

To estimate the pleiotropic effects of variants across traits, we also performed a meta-analysis of SNP effects estimated using LMMs with an empirical Bayesian multivariate adaptive shrinkage approach that included results from nine traits [70]. Over 150 variants exhibited strong associations across the nine traits with most associations occurring in chromosomes six and seven (Figure S14). While most associations occur within known loci including *Dw3*, *Ma3*, *Ma6*, and the *D* locus, a SNP on Chr10 colocalizes with previously identified locus for fresh stem weight and juice yield [97] as well as sucrose content [82]. Taken together, these multivariate approaches highlight the pleiotropic effects of loci across the sorghum genome and support the importance of collecting peripherally related phenotypes to maximize carbon accumulation and partitioning.

## Discussion

The diverse carbon-partitioning regimes of sorghum have the potential to provide valuable insights into the genetic control of carbon partitioning in grasses [98] from transport [38] to compartmentalization [99]. Genes sensitive to carbohydrate concentration compose part of a highly conserved network necessary for cellular adjustment to nutrient availability and the partitioning of carbon among tissues and organs [100]. A holistic understanding of these processes requires multiscale phenotypes from molecule-specific quantification to anatomically aggregated measures of carbon. These multiscale metrics are necessary to accurately assess traits such as biomass where optical measures are typically poorly correlated with manually collected, macroscale phenotypes [3]. Orthogonal and partially correlated measures of diverse morphology assist in resolving functional questions of plant growth and development while simultaneously improving significant associations with functional genomic data. In conjunction with broad-scale phenotyping, multiparameter statistical approaches improve inferences through joint consideration of genomic and phenotypic measures [3].

Here, we identified numerous putative loci that are associated with a variety of different phenotypes across different scales. The interactions between loci and locus pleiotropy hint at the underlying genetic architecture of these dynamic carbon-partitioning traits. However, source and sink interactions can complicate the dissection of individual traits [33]. High-capacity, non-photosynthetic sinks can increase yield through sugar-responsive genes that mediate feed-forward loops that ultimately bolster systemic carbon accumulation [37, 101]. Conversely, as seen here with the *D* locus NAC transcription factor exhibiting reduced lignin content but increased sugar and grain yields [84], selection for some loci can result in a tradeoff between carbon regimes. The identification of these feedback mechanisms suggests that ongoing optimization of carbon allocation should simultaneously focus on improved source and sink strengths as a system of interconnected processes from nitrogen assimilation to photosynthetic efficiency [29, 33, 37]. Additionally, these findings indicate that sorghum yields (i.e., sugar, grain, forage, and biomass) may be further optimized to incorporate beneficial alleles from other sorghum types.

Using these models, we identified numerous candidate loci associated with carbon-partitioning traits using the CP-NAM. Several traits, such as WSC and biomass traits, shared associated loci supporting previous observations that selection for non-target, sink-related traits may collectively increase yields across carbon-partitioning regimes. Future studies should examine multi-trait, multi-environment data to further extricate the environmental and genotype-by-environment effects on carbon-partitioning traits by leveraging the power of multiscale traits and MV-LMMs [102]. Additionally, breeders may consider collecting peripherally related traits as a means of understanding and maximizing carbon flow in their system as selection for carbon sinks is not a zero-sum relationship.

## Supporting information

NAM_Plot_data

Blink_results

Heritabilities

Bayes_factors

MLM_results

NAM_compositional_data

QTL_results

## Declarations

### Authors’ contributions

JLB wrote the manuscript. JLB performed all computational analyses. AC, MM, NK, and SS managed field experiments. AC, KEJ, MM, NK, and SS collected phenotypes. JLB, SS, and SK conceptualized, developed, and implemented the study design.

### Availability of data and material

Supplemental figures and tables are available with submission.

### Code availability

Scripts are available on GitHub (https://github.com/jlboat/CP-NAM_2021) under MIT License.

### Consent for publication

All authors reviewed and approved the final manuscript.

## Acknowledgments

Computational analyses were performed on Clemson University’s Palmetto Cluster, and we thank the staff who assisted with cluster and software maintenance.

