## Supplemental_figures for "Dissecting the Genetic Architecture of Carbon Partitioning in Sorghum using Multiscale Phenotypes"

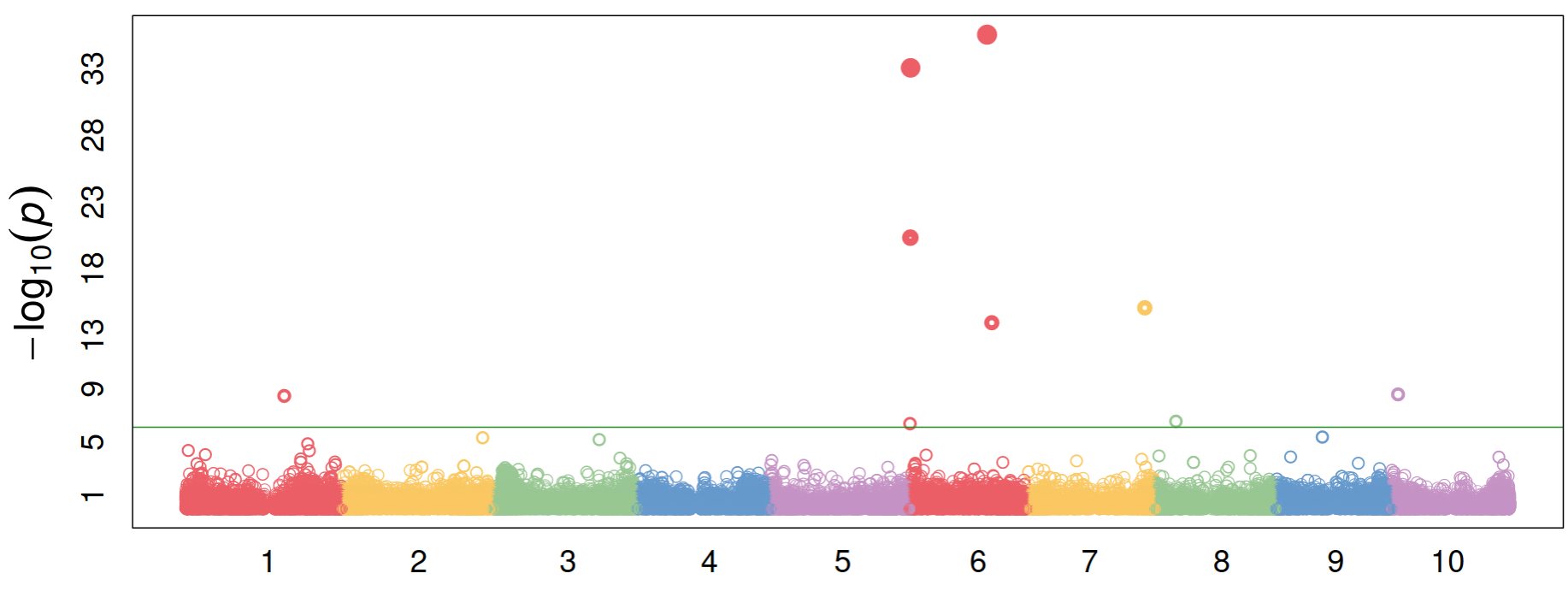

**Figure S1**. Manhattan plot of Blink results for plant height.

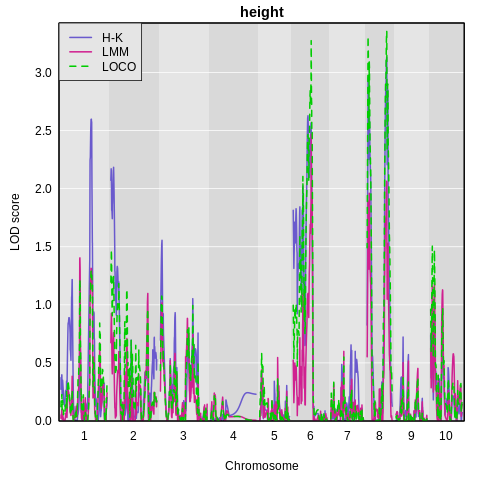

**Figure S2.** QTL mapping for plant height in PI586454 using Haley-Knott (H-K), linear mixed model (LMM), and leave-one-chromosome-out (LOCO) methods.

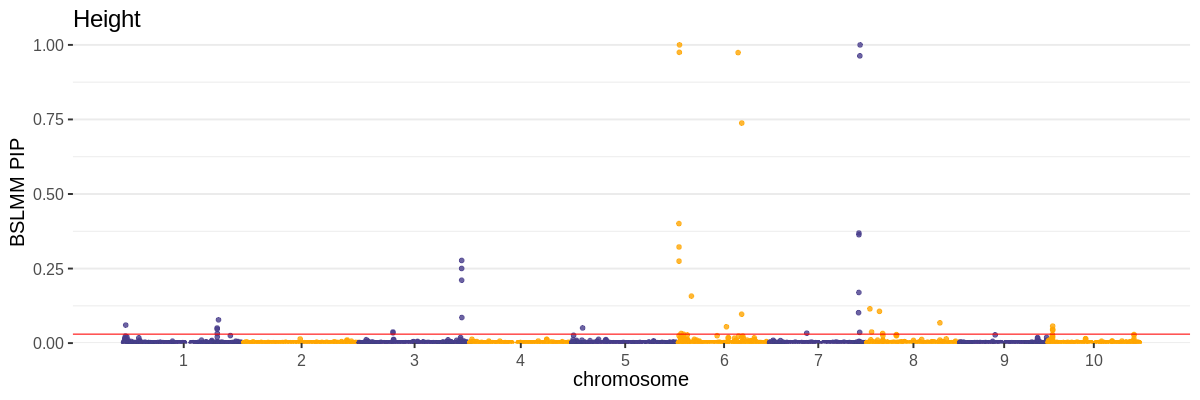

**Figure S3**. Manhattan plot of Bayesian sparse linear mixed model posterior inclusion probability (PIP) for SNPs across the sorghum genome generated using GEMMA. The horizontal, red line indicates the significance threshold for SNPs (0.036).

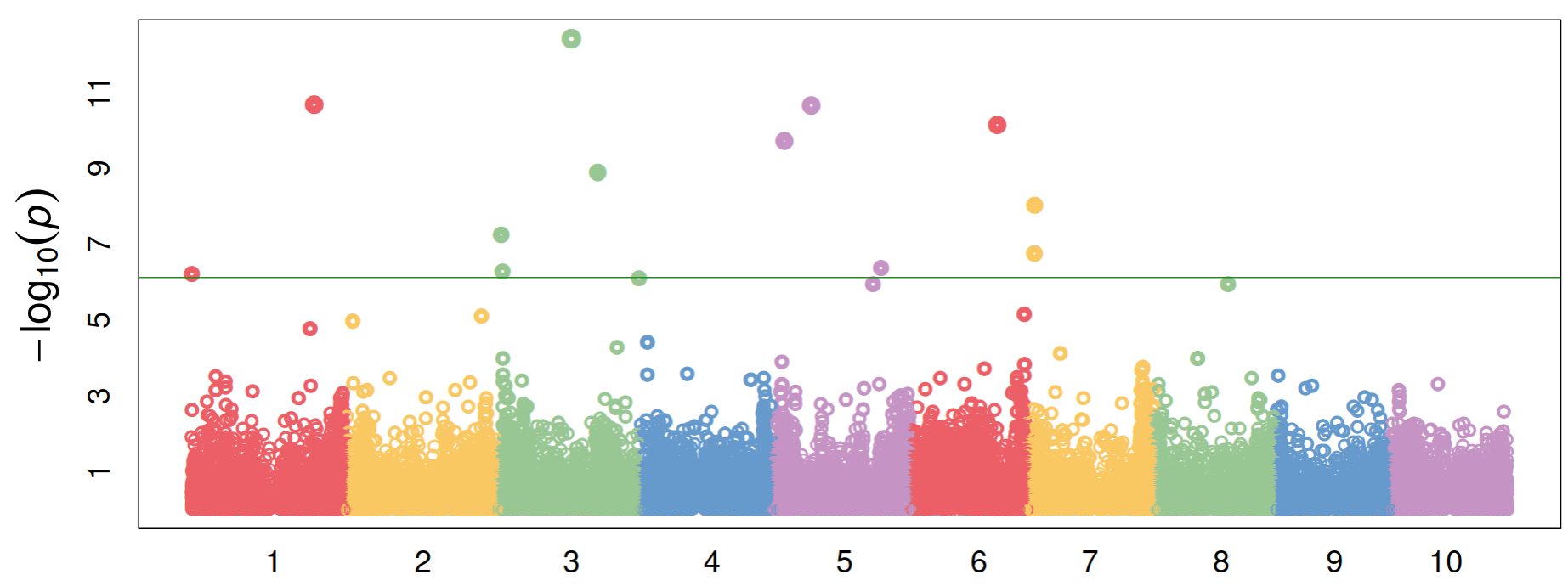

**Figure S4.** Manhattan plot of Blink results for days to harvest.

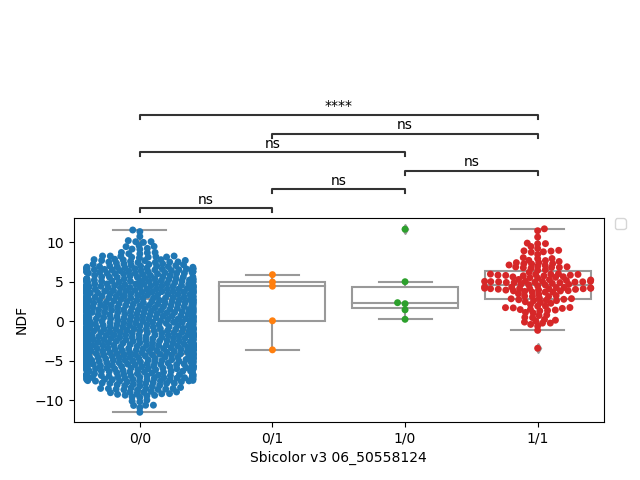

**Figure S5**. Boxplots represent the phenotypic distribution for NDF across genotypes. Pairwise t-tests were performed between genotypes as indicated on top of the figure where ns means not significant and **** indicates a p-value <= 1.00e-04.

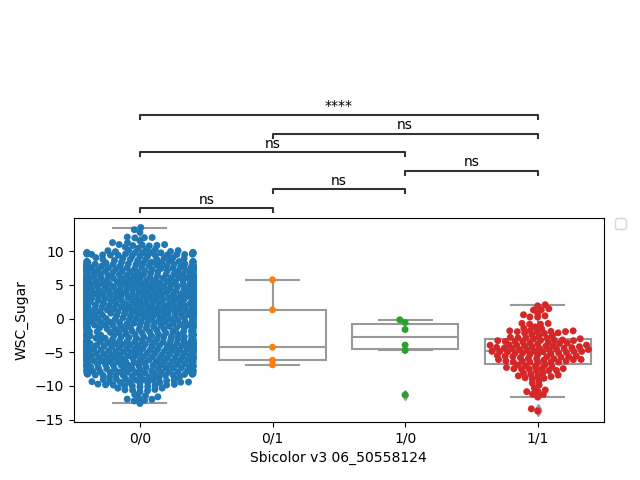

**Figure S6**. Boxplots represent the phenotypic distribution for WSC across genotypes. Pairwise t-tests were performed between genotypes as indicated on top of the figure where ns means not significant and **** indicates a p-value <= 1.00e-04.

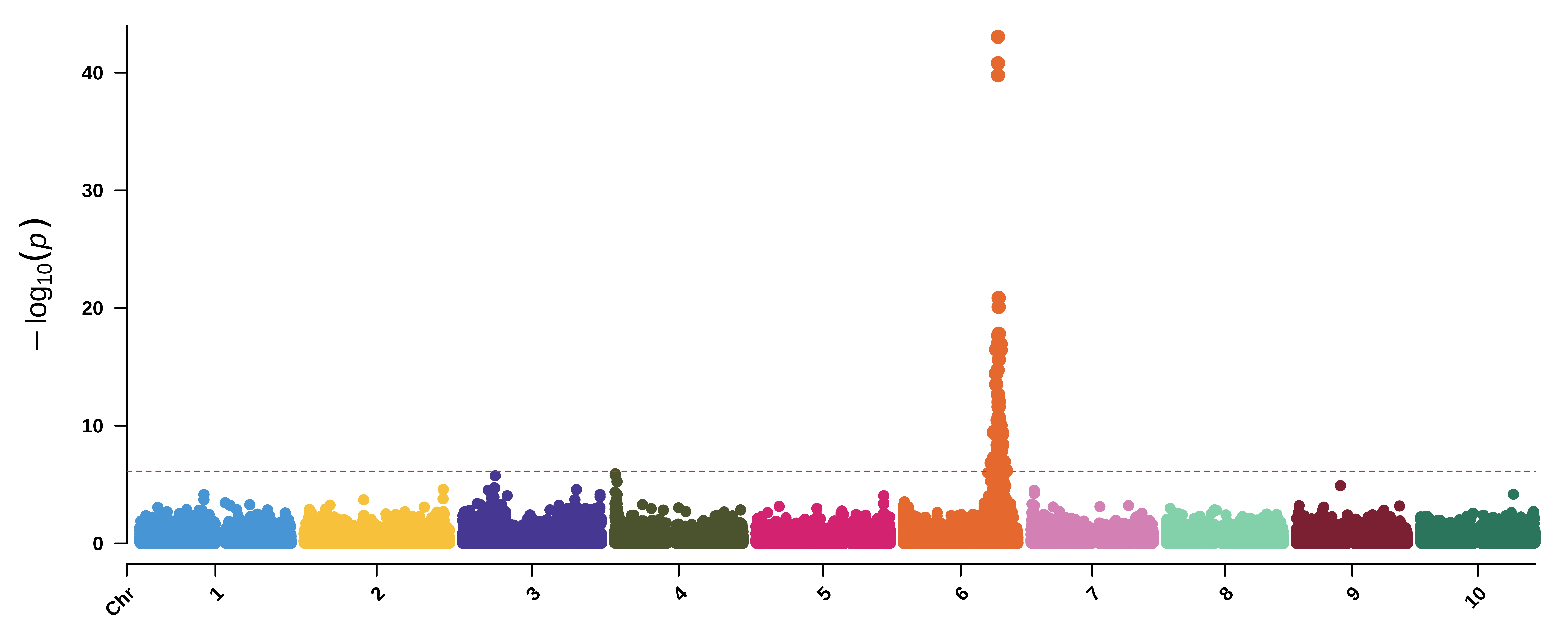

**Figure S7**. Manhattan plot of a MV-LMM using GEMMA with WSC and NDF as response variables and maturity and DTH as covariates. The red-dashed, horizontal line represents the Bonferroni-corrected significance threshold.

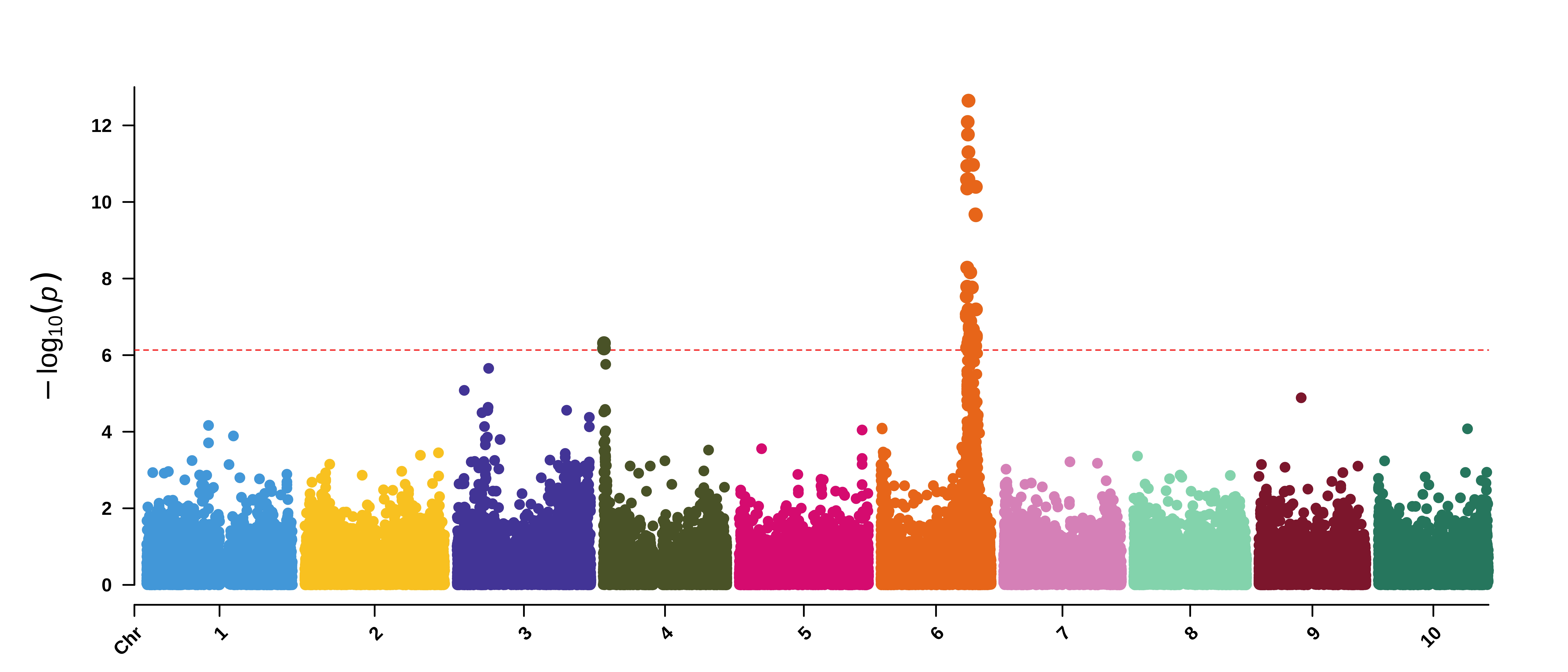

**Figure S8**. Manhattan plot of a MV-LMM using GEMMA with WSC and NDF as response variables and maturity, DTH and the top SNP from the MV-LMM represented in **Figure S7** as covariates. The red-dashed, horizontal line represents the Bonferroni-corrected significance threshold.

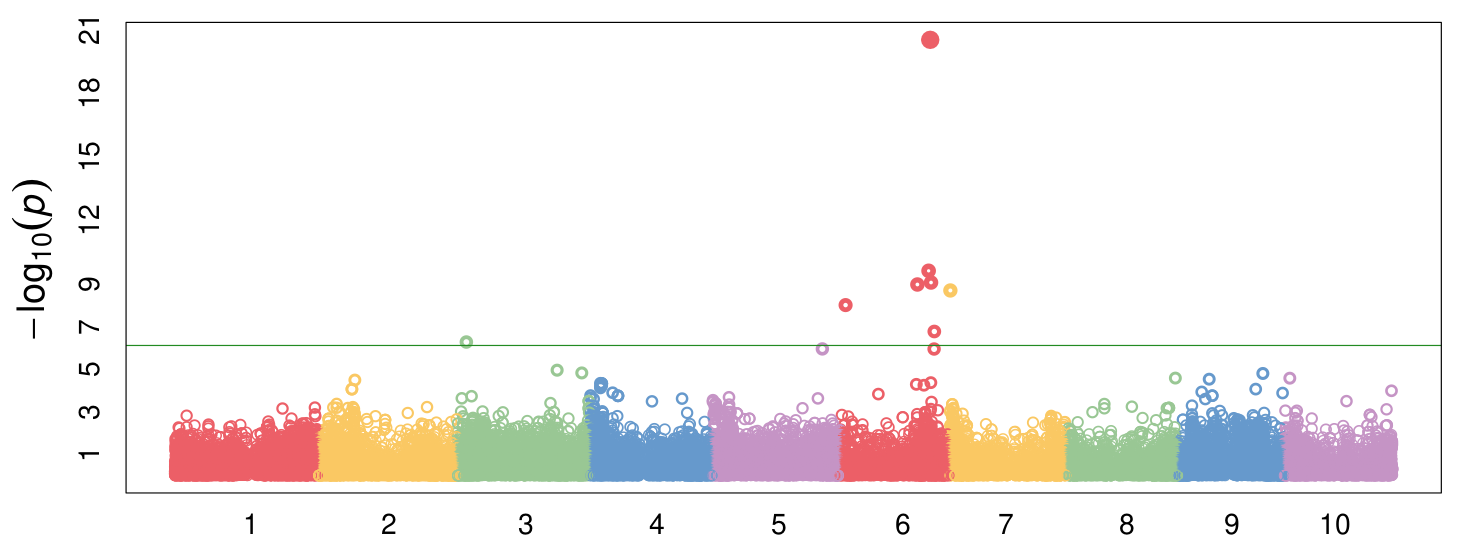

**Figure S9**. Manhattan plot of Blink results for ADF with maturity and DTH covariates.

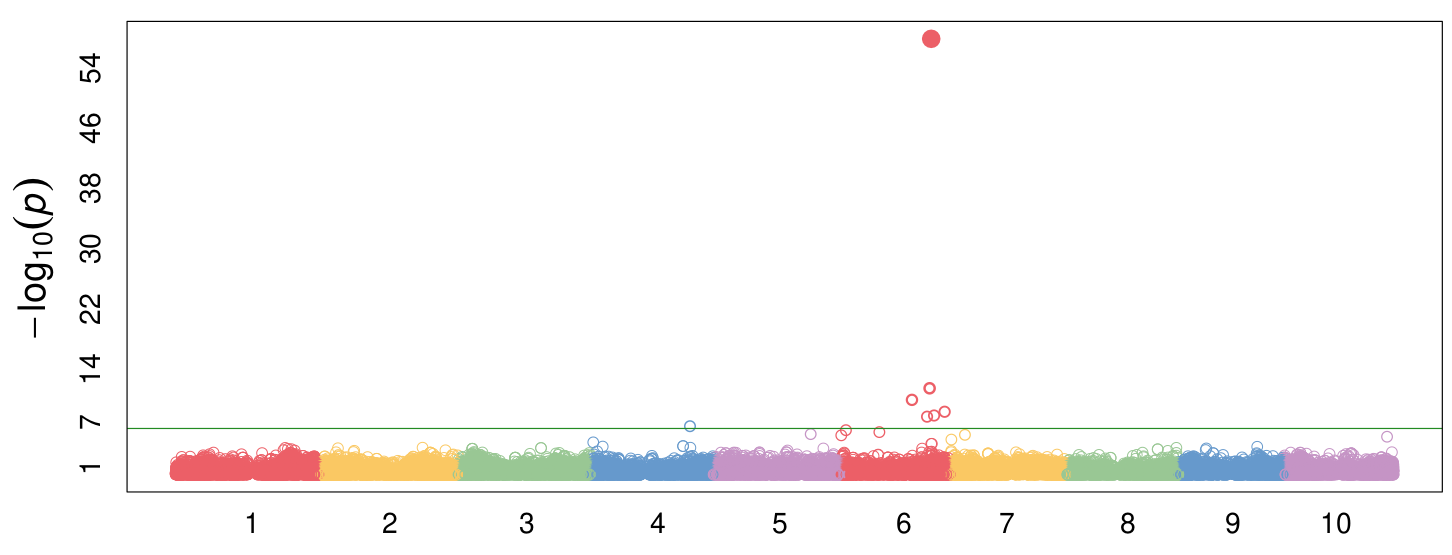

**Figure S10**. Manhattan plot of Blink results for NDF with maturity and DTH covariates.

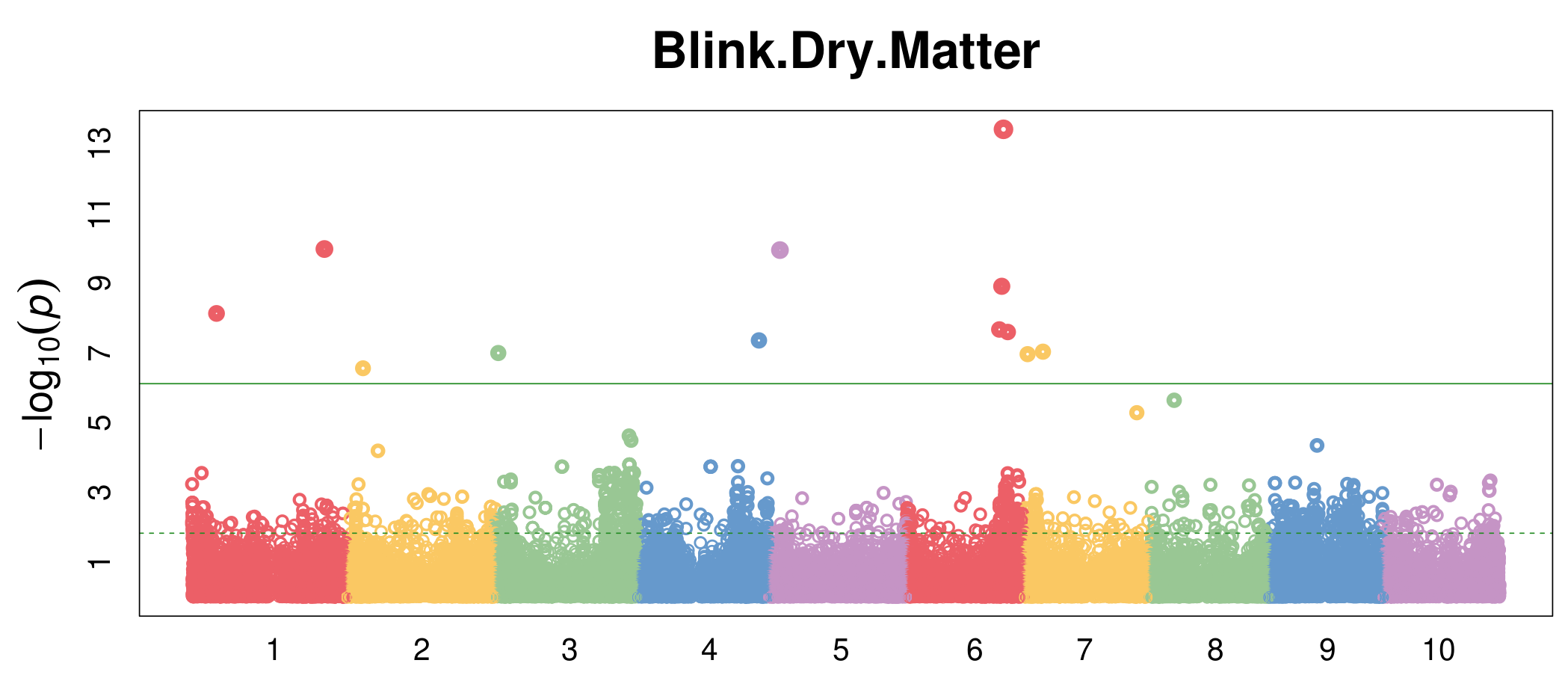

**Figure S11**. Dry matter Manhattan plot of Blink results. The block around Chr04:63,328,486 spans up to 65,734,119 thereby encompassing the VIT gene.

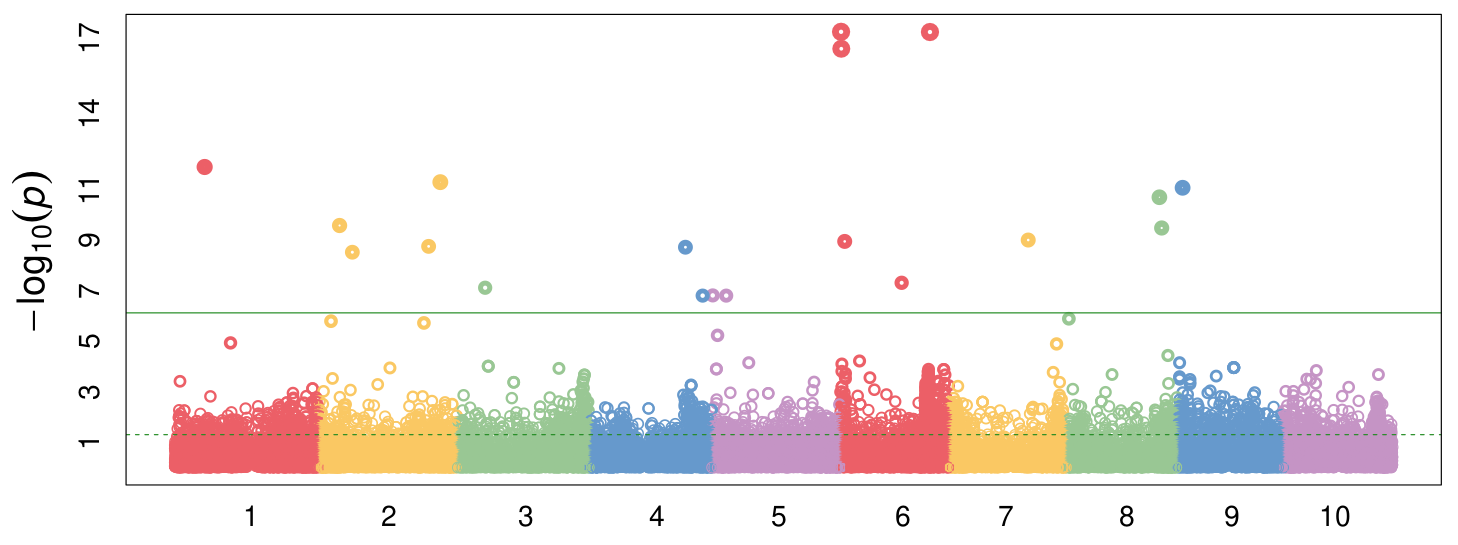

**Figure S12**. Manhattan plot of Blink results for adjusted crude protein.

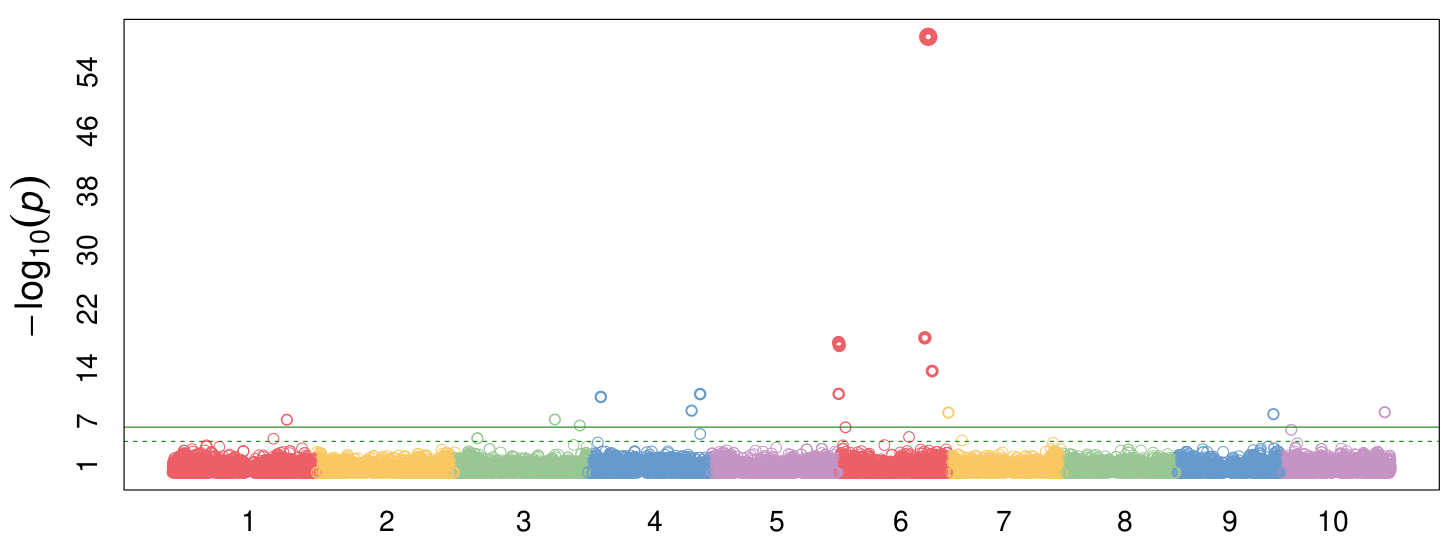

**Figure S13**. Manhattan plot of Blink results for NEG.

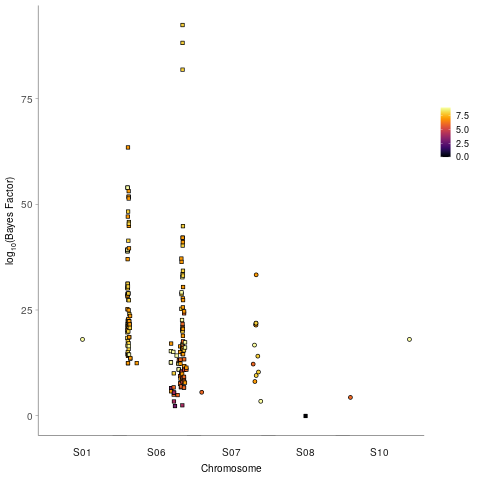

**Figure S14**. Manhattan plot of the Bayes Factor estimates from mashr for LMMs for ADF, ash, dry matter, dry weight, height, NDF, P, wet weight, and WSC.
